# New alleles of *D-2-hydroxyglutarate dehydrogenase* enable studies of oncometabolite function in *Drosophila melanogaster*

**DOI:** 10.1101/2025.03.27.645621

**Authors:** Madhulika Rai, Prince Okah, Shefali A. Shefali, Alexander J. Fitt, Michael Z. Shen, Mandkhai Molomjamts, Robert Pepin, Travis Nemkov, Angelo D’Alessandro, Jason M. Tennessen

**Affiliations:** Department of Biology, Indiana University, Bloomington, IN 47405, USA; Department of Chemistry, Indiana University, Bloomington, IN 47405, USA; Department of Biochemistry and Molecular Genetics, University of Colorado Anschutz Medical Campus, Colorado, USA; Affiliate Member, Melvin and Bren Simon Cancer Center, Indianapolis, IN, 46202, USA

**Keywords:** *Drosophila melanogaster*, metabolism, D-2HG, oncometabolite, d-2-hydroxyglutarate dehydrogenase, D2hgdh

## Abstract

D-2-hydroxyglutarate (D-2HG) is a potent oncometabolite capable of disrupting chromatin architecture, altering metabolism, and promoting cellular dedifferentiation. As a result, ectopic D-2HG accumulation induces neurometabolic disorders and promotes progression of multiple cancers. However, the disease-associated effects of ectopic D-2HG accumulation are dependent on genetic context. Specifically, neomorphic mutations in the mammalian genes *Isocitrate dehydrogenase 1* (*IDH1*) and *IDH2* result in the production of enzymes that inappropriately generate D-2HG from α-ketoglutarate (αKG). Within this genetic background, D-2HG acts as an oncometabolite and is associated with multiple cancers, including several diffuse gliomas. In contrast, loss-of-function mutations in the gene *D-2-hydroxyglutarate dehydrogenase* (D2hgdh) render cells unable to degrade D-2HG, resulting in excessive buildup of this molecule. *D2hgdh* mutations, however, are not generally associated with elevated cancer risk. This discrepancy raises the question as to why ectopic D-2HG accumulation in humans induces context-dependent disease outcomes. To enable such genetic studies *in vivo*, we generated two novel loss-of-function mutations in the *Drosophila melanogaster* gene *D2hgdh* and validated that these alleles result in ectopic D-2HG. Moreover, we observed that *D2hgdh* mutations induce developmental and metabolomic phenotypes indicative of elevated D-2HG accumulation. Overall, our efforts provide the *Drosophila* community with new mutant strains that can be used to study D-2HG function in human disease models as well as in the context of normal growth, metabolism, and physiology.

## INTRODUCTION

D-2-hydroxyglutarate (D-2HG) is a hydroxy acid produced from the citric acid cycle intermediate alpha-ketoglutarate (αKG) (YE *et al*. 2018; DU AND HU 2021). Under physiological conditions, animals can synthesize D-2HG by two enzymatic mechanisms: (i) the enzyme hydroxyacid-oxoacid transhydrogenase (HOT) interconverts αKG and γ-hydroxybutyrate into succinic semialdehyde and D-2HG (STRUYS *et al*. 2005b); and (ii) the noncanonical activity of the enzyme d-3-phosphoglycerate dehydrogenase reduces αKG to form D-2HG (FAN *et al*. 2015). However, despite being a side-product of normal cellular metabolism, intracellular D-2HG concentration remain relatively low due, in part, to the activity of the mitochondrial enzyme D-2-hydroxyglutarate dehydrogenase (D2HGDH), which converts D-2HG into αKG in an FAD dependent manner (ACHOURI *et al*. 2004).

The D2hgdh-imposed limitation on D-2HG pool size is essential for normal cellular function, as D-2HG is a potent signaling molecule capable of inhibiting a wide-range of αKG-dependent dioxygenases, including the Jmj class of histone lysine demethylases, the Tet family of enzymes, and prolyl hydroxylases (FIGUEROA *et al*. 2010; CHOWDHURY *et al*. 2011). As a result, increased D-2HG accumulation can induce dramatic changes in chromatin architecture, gene expression, and cellular metabolism, and ultimately lead to a variety of disease phenotypes (YE *et al*. 2018; DU AND HU 2021). For example, loss-of-function mutations in the human *D2hgdh* gene result in an inborn error of metabolism known as D-2-hydroxyglutaric aciduria, which is characterized by elevated urinary D-2HG levels, developmental delays, seizures, and brain abnormalities (STRUYS *et al*. 2005a; STRUYS 2006). Similarly, conditional deletion of *D2hgdh* in the mouse brain results in increased D-2HG accumulation and leads to impaired activation of radial-glia-like neural stem cells, reduced histone acetylation, and impairment of hippocampus-dependent learning (LIU *et al*. 2023). Together, these findings highlight the essential role of D2hgdh in limiting pathological D-2HG accumulation.

Because D-2HG interferes with αKG-dependent processes, this molecule is also capable of acting as an oncometabolite and is well-documented to promote the growth and development of a wide range of tumors (YE *et al*. 2018; WANG *et al*. 2020; CHOU *et al*. 2021; DU AND HU 2021). The oncogenic activity of this metabolite, however, is sensitive to the genetic mechanism responsible for inducing aberrant D-2HG accumulation. In nearly all documented studies of D-2HG-positive human tumors, the underlying causes are neomorphic gain-of-function mutations that affect either arginine 132 of IDH1 or arginine 172 of IDH2 (DANG *et al*. 2009; LOSMAN AND KAELIN 2013). These mutations alter the catalytic activity of IDH1/IDH2 in a manner that induces inappropriate D-2HG production (LOSMAN AND KAELIN 2013). Such mutations are found in a wide variety of human tumors and are particularly common in lower-grade gliomas, with 70%-80% of patients with grade II and III astrocytomas and oligodendrogliomas harboring *IDH1* mutations (YAN *et al*. 2009). Similar findings have emerged from mice, where expression of IDH1(R132H) in astrocytes induces D-2HG production and promotes glioma development in sensitized backgrounds (PHILIP *et al*. 2018). In fact, the widespread occurrence of *IDH1/IDH2* mutations in gliomas and acute myeloid leukemia has led to the successful development of FDA approved IDH inhibitors (GATTO *et al*. 2021; TIAN *et al*. 2022), which are now used in combination therapies to treat IDH1/IDH2-positive tumors.

While there is overwhelming evidence that D-2HG acts as an oncometabolite in cells that express mutant IDH1/IDH2, there is limited evidence indicating that changes in D2hgdh expression or activity drive glioma progression, with only a handful of studies hinting this possibility (LIN *et al*. 2015; ECKEL-PASSOW *et al*. 2020). The paucity of evidence linking *D2hgdh* mutations with glioma occurrence is striking considering the wide-spread prevalence of *IDH1/IDH2* mutations in these tumors (YAN *et al*. 2009). Such observations raise the question as to why D-2HG only acts as an oncometabolite in specific genetic backgrounds and highlights the need to further study endogenous D-2HG metabolism in both healthy tissues and disease models. Towards this goal, we are using the fruit fly *Drosophila melanogaster* as a model to understand how loss of D2hgdh activity affects metabolism, growth, and development.

Similar to humans and mice, flies ubiquitously expressing an oncogenic *Idh* mutant allele (*UAS-Idh-R195H)* accumulate elevated D-2HG levels. Moreover, *UAS-Idh-R195H* expression in blood cells using *hml-Gal4* induces melanotic mass formation and excess hemocyte production (REITMAN *et al*. 2015). This finding demonstrates that expression of a neomorphic Idh variant in flies partially phenocopies the disease associated Idh1/2 phenotypes observed in other organisms. Furthermore, overexpression of D2hgdh from a *UAS-D2hgdh* transgene reduces D-2HG accumulation in flies expressing *UAS-Idh-R195H*, indicating that *Drosophila* D2hgdh also functions to degrade D-2HG (REITMAN *et al*. 2015). However, these earlier *Drosophila* studies primarily focused on the oncogenic effects of mutant Idh expression and largely overlooked the endogenous biological roles of D-2HG and D2hgdh. We would also note that while several studies focusing on L-2HG have also quantified D-2HG levels (LI *et al*. 2017; LI *et al*. 2018; MAHMOUDZADEH *et al*. 2020), these investigations did not examine the role of D-2HG in development, physiology, or metabolism.

To facilitate additional studies of D-2HG metabolism in *Drosophila*, we used CRISPR/Cas9 approach to generate loss-of-function mutations in *D2hgdh*. Here we demonstrate that the resulting mutant lines are viable and display elevated D-2HG levels. Moreover, *D2hgdh* mutants exhibit decreased fecundity and viability, as well as occasional formation of melanotic masses. Finally, metabolomic analysis of adult male *D2hgdh* mutants reveal significant changes in essential amino acids levels. Overall, our findings establish *Drosophila* as a powerful genetic model for studying endogenous D-2HG metabolism and suggest that future studies of the *D2hgdh* mutants will advance our understanding of how loss of D2hgdh activity results in disease phenotypes.

## MATERIALS AND METHODS

### Drosophila melanogaster husbandry

Fly stocks were maintained at 25°C on Bloomington Drosophila Stock Center (BDSC) food. Larvae were raised on molasses agar plates with yeast paste spread over the surface as previously described (LI AND TENNESSEN 2017). Since all mutations were generated in a *w^1118^*background, we used *w^1118^* as a control for all experiments. For all adult analysis, males and virgin females were collected within 8 hours of eclosion, the sexes separated into vials containing BDSC food, and animals were aged for 5-7 days. Flies were then anesthetized using carbon dioxide and collected immediately for subsequent analysis. Flybase was used as an essential reference tool throughout this study (GRAMATES *et al*. 2022; ÖZTÜRK-ÇOLAK *et al*. 2024).

### Generation of *D2hgdh* mutations

*D2hgdh* mutations were generated using CRISPR/Cas9 (GRATZ *et al*. 2013; SEBO *et al*. 2014). An oligo encoding a guide RNA sequence that targeted exon 3 of the *D2hgdh* locus (5’ CTTCGCTTAAGCCCGGAAGCACGG 3’) was inserted into the BbsI site of pU6-BbSI-gRNA (Addgene). The gRNA construct was injected into RRID:BDSC_51324 (w[1118]; PBac{y[+mDint2]=vas-Cas9}VK00027). Two mutations isolated from independently injected embryos were identified by PCR amplification of the region surrounding exon 2 of the *D2hgdh* locus using oligos 5’-TCCTCCACGATGAGATTCCAAC -3’ and 5’-TGATCGAAGTTCTCCAGGATGC -3’. The resulting PCR product was sequenced using oligos 5’-CCTATCACCACCACCACCATC-3’ and 5’-AAGGACAATCTCGTCGCAGATC-3’. This approach identified two *D2hgdh* deletions: *D2hghd^12-6^* consists of a 76 bp deletion (5’-CCACTCTGACCGACAAGGATGTGGCGCATTTCGAGCAGCTCCTGGGCAAGAACTTCGTGCTCACTGAGGACCTGGA-3’). *D2hghd^5-5^* contains a 5 bp deletion of 5’-GACCT-3’ within the sequence 5’-GAACTTCGTGCTCACTGAG<u>GACCT</u>GG-3’ (see Figure 1A).

**Figure 1.**
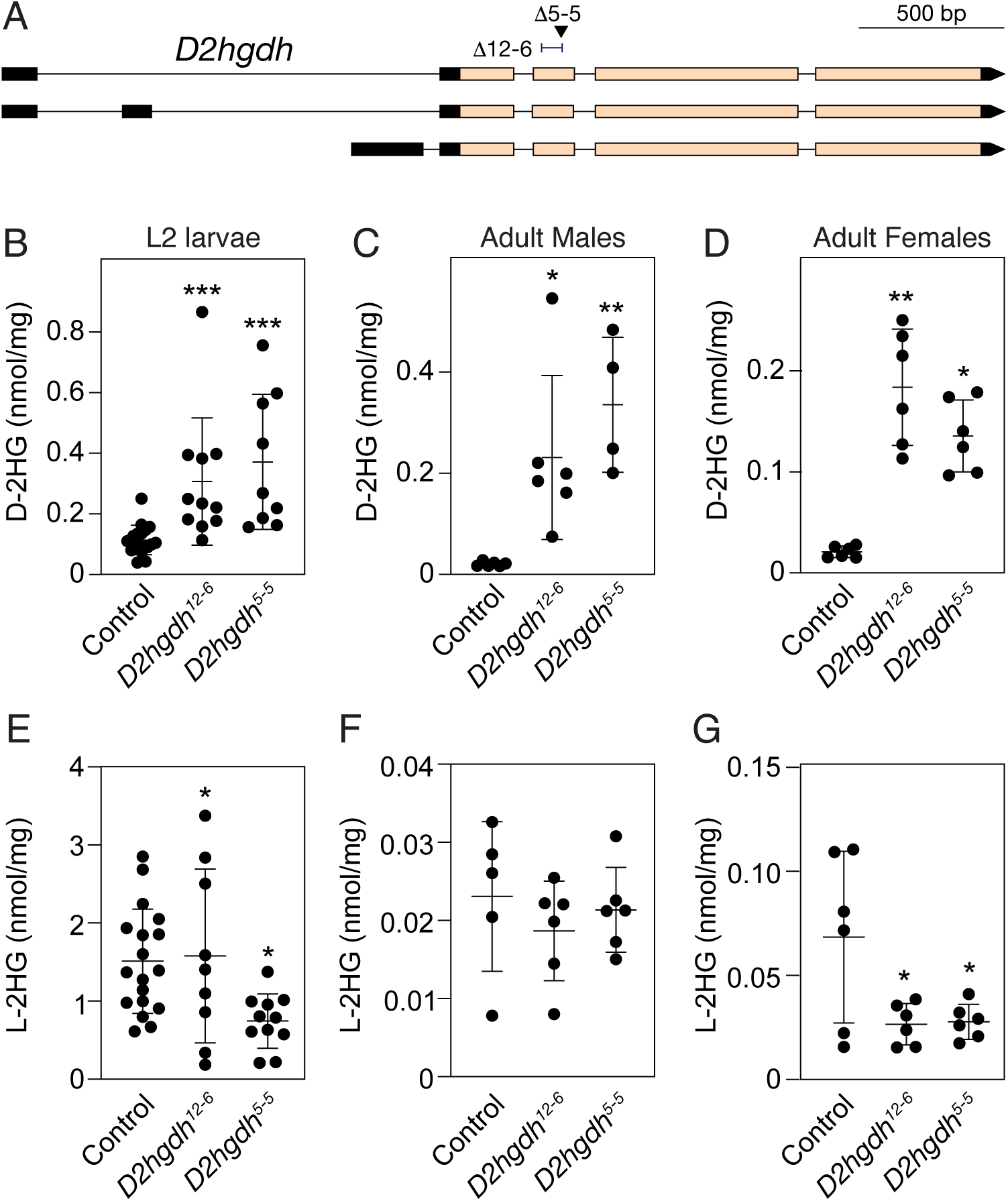
*D2hgdh* mutations induce elevated D-2HG accumulation. (A) A schematic diagram illustrating the *D2hgdh* locus and the mutations *D2hgdh^12-6^* and *D2hgdh^5-5^*. The arrow under Δ5-5 points to the location of the *D2hgdh^5-5^* 7 bp deletion. The bracket next to Δ12-6 illustrates to the location of the *D2hgdh^5-5^* 76 bp deletion. (B-G) The amount of D-2HG and L-2HG were simultaneously quantified in both control and *D2hgdh* mutants during the (B,E) L2 larval stage as well as in (C,F) adult males and (D,G) adult females. Data presented as a scatter plot with the lines representing the mean and standard deviation. *P*-values calculated using a Wilcoxon/Kruskal-Wallis tests followed by a Dunn’s test to compare mutant sample with control. \**P*<0.05. \*\**P*<0.01. \*\*\**P*<0.001. A minimum of six biological replicates were analyzed for each genotype. Each replicate contained either 25 mid-L2 larvae or 20 adults (5-7 days post-eclosion). Experiments were repeated three times.

### Gas Chromatography-Mass Spectrometry (GC-MS) analysis

D- and L-2HG were quantified using a protocol specifically designed for analyzing these two metabolites (LI AND TENNESSEN 2019). Briefly, either 25 mid-L2 larvae or 20 adults (5-7 days post-eclosion) were placed in a pre-tared 2 ml screw cap tube containing 1.4 mm ceramic beads (Catalog No. 15-340-153; Fisher Scientific), the mass was determined with an analytical balance, and the tube was immediately dropped into liquid nitrogen and subsequently stored at - 80°C until processing. 800 μl of ice cold extraction buffer (90% MeOH with 8 mg of 2,2,3-d3-R,S-2-hydroxyglutarate) was then added to each sample (kept in -20°C enzyme caddies) and tubes were then immediately placed in a Omni Bead Ruptor 24 and homogenized for 30 seconds at 6.4 m/s. Samples were subsequently incubated at -20°C for 1 hr. After the incubation, samples were centrifuged at 20,000 x g for 5 min at 4°C. 600 μl of the supernatant was then transferred into a 1.5 mL microfuge tube and dried overnight using a vacuum centrifuge.

Dried samples were resuspended in 50 μl of R-2-Butanol and 5 μl of HCL and incubated at 90°C for three hours with shaking at 300 rpm using an Eppendorf ThermoMixer F1.5. After cooling, 200 μl of water and 500 μl of hexane was added to each tube. The organic phase (hexane) was then transferred to a new tube and dried for 30 minutes using a vacuum centrifuge. The dried samples were resuspended in 60 μl of acetic anhydride and 60 μl of pyridine and incubated at 80°C for 1 hr with shaking at 300 rpm. Samples were then dried for 3 hrs using a vacuum centrifuge and resuspended in 60 μl of hexane. GC-MS analyses were performed using an Agilent GC6890-5973i MS equipped with a Gerstel MPS autosampler and a 30 m Phenomex ZB5-5 MSi column.

### Ultra High-pressure Liquid Chromatography - Mass Spectrometry (UHPLC-MS)-based Metabolomics

*D2hgdh* mutants and the control strain were raised on BDSC food at 25°C and groups of 20 newly eclosed adult males were transferred to a fresh vial and aged for 5 days, at which point they were placed in a 1.5 mL microfuge tube and drop frozen in liquid nitrogen. Samples were then individually transferred to a pretared 1.4 mm bead tube, the mass was measured using a Mettler Toledo XS105 balance, and the sample was returned to a liquid nitrogen bath prior to being stored at -80°C. Samples were subsequently homogenized in 0.8 mL of prechilled (-20 °C) 90% methanol containing 2 μg/mL succinic-d4 acid, for 30 seconds at 6.45 m/s using a bead mill homogenizer located in a 4°C temperature control room. The homogenized samples were incubated at -20 °C for 1 hour and then centrifuged at 20,000 × g for 5 min at 4°C. The resulting supernatant was sent for metabolomics analysis at the University of Colorado Anschutz Medical Campus.

UHPLC-MS metabolomics analyses were performed as previously described (NEMKOV *et al*. 2019). Briefly, the analytical platform employs a Vanquish UHPLC system (Thermo Fisher Scientific, San Jose, CA, USA) coupled online to a Q Exactive mass spectrometer (Thermo Fisher Scientific, San Jose, CA, USA). The (semi)polar extracts were resolved over a Kinetex C18 column, 2.1 x 150 mm, 1.7 µm particle size (Phenomenex, Torrance, CA, USA) equipped with a guard column (SecurityGuard^TM^ Ultracartridge – UHPLC C18 for 2.1 mm ID Columns – AJO-8782 – Phenomenex, Torrance, CA, USA) using an aqueous phase (A) of water and 0.1% formic acid and a mobile phase (B) of acetonitrile and 0.1% formic acid for positive ion polarity mode, and an aqueous phase (A) of water:acetonitrile (95:5) with 1 mM ammonium acetate and a mobile phase (B) of acetonitrile:water (95:5) with 1 mM ammonium acetate for negative ion polarity mode. The Q Exactive mass spectrometer (Thermo Fisher Scientific, San Jose, CA, USA) was operated independently in positive or negative ion mode, scanning in Full MS mode (2 μscans) from 60 to 900 m/z at 70,000 resolution, with 4 kV spray voltage, 45 sheath gas, 15 auxiliary gas. Calibration was performed prior to analysis using the Pierce^TM^ Positive and Negative Ion Calibration Solutions (Thermo Fisher Scientific).

### Statistical Analysis of Metabolomics Data

Both metabolomics datasets were analyzed using Metaboanalyst 5.0 (PANG *et al*. 2021). Samples were normalized to mass and the data was preprocessed using log normalization and Pareto scaling. Preprocessed data can be found in Table S1. The Quantitative Enrichment analysis tool was used to conduct the enrichment analysis in Figures S1 and S2.

### Characterization of Developmental Phenotypes

Twenty-five males and 50 virgin females for the control and mutant genotypes were placed in an egg laying chamber and placed in a 25°C incubator. For fecundity assays, flies were mated for 24 hours prior to start of the assay, after which egg-laying plates were placed in the chamber for one-hour intervals and the number of eggs were counted by visual observation in using a standard dissection microscope. Embryonic viability was measured by counting the number of eggs on each egg-laying plate that hatched 30 hrs after egg-laying. Pupal viability was measured by transferring 20 newly hatched L1 larvae to a food vial and subsequently counting the number of pupae until 8 days after egg-laying.

### Statistical Analysis

All data analyses were conducted using Graphpad Prism 9. *P*-values were calculated using a Wilcoxon/Kruskal-Wallis tests followed by a Dunn’s test to compare mutant sample with control.

## RESULTS

### *D2hgdh* mutants accumulate excess D-2HG

As a first step towards determining how D2hgdh activity regulates D-2HG levels during the *Drosophila* life-cycle, we used CRISPR/Cas9 to generate two mutations in the *D2hgdh* gene. As described in the methods, our approach isolated two *D2hgdh* mutant alleles consisting of a 5 bp deletion (*D2hgdh^5-5^*) and a 76 bp deletion (*D2hghd^12-6^*), both of which disrupt the conserved catalytic region of the enzyme (Figure 1A; see Methods for a description of the deleted regions). The resulting mutants are homozygous/hemizygous viable, thus allowing us to determine if *D2hgdh* mutations alter steady state D-2HG levels at multiple points during the life-cycle. Consistent with the conserved role of D2hgdh in catabolizing D-2HG, both *D2hgdh^5-5^*and *D2hghd^12-6^* mutant males, females, and L2 larvae displayed significantly higher D-2HG levels as compared with controls (Figure 1B-D). In contrast, L-2HG levels were either unchanged or decreased in *D2hgdh* mutants, indicating that D2hgdh activity is specific to D-2HG (Figure 1E-G). These results demonstrate that *D2hgdh* encodes the enzyme responsible for limiting D-2HG accumulation in the fly.

### *D2hgdh* mutants develop melanotic masses and display developmental defects

*D2hgdh* mutations in humans lead to developmental delays and as well as hypermelanization of the abdominal region – a condition termed angiokeratoma, which is characterized by benign skin lesions (PRESTON *et al*. 2019). Consistent with these human disease phenotypes, we observe similar phenomenon in *D2hgdh* mutants - both *D2hgdh^5-5^* and *D2hghd^12-6^* mutant larvae exhibit melanotic masses within their abdominal cavity (Figure 2A-C). While these lesions are relatively rare (present in ∼2% of larvae; Figure 2C), the masses are easily observed and occur at similar rates in both strains. Moreover, a closer examination of the lesions using confocal microscopy reveal that these masses were often associated with regions of the hindgut that lacked nuclei (Figure 2D-I). While the significance of these lesions remains unclear, we note that a similar observation was reported in larvae expressing the *UAS-Idh-R195H* transgene in hemocytes using *hml-Gal4* (REITMAN *et al*. 2015), suggesting that these masses stem from elevated D-2HG levels during blood cell development.

**Figure 2.**
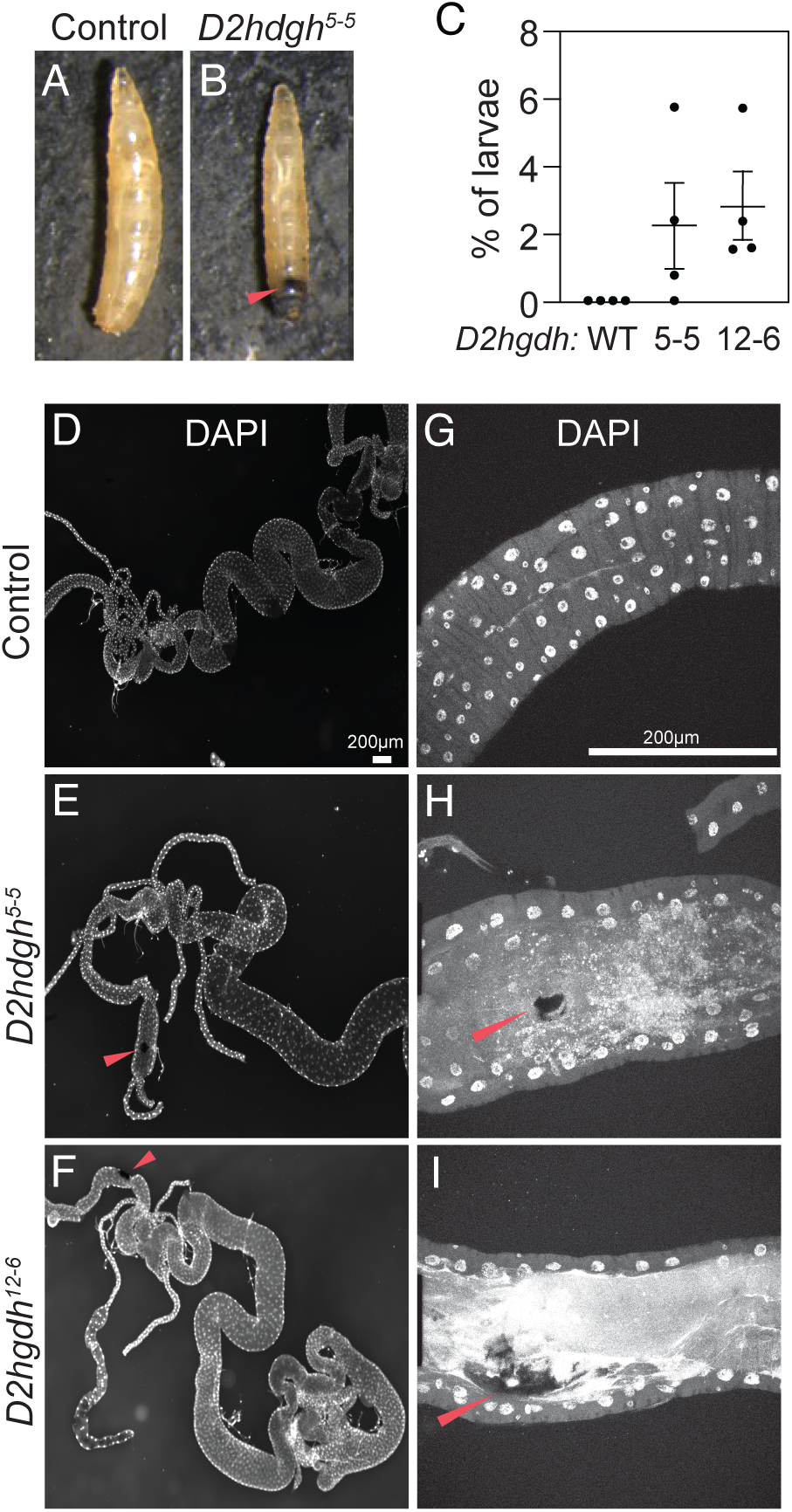
*D2hgdh* mutant exhibit melanotic mass formation. Unlike (A) *w^1118^* controls, (B) *D2hgdh* mutant develop melanotic masses in their abdominal region. (C) The melanotic mass phenotype occurs in ∼2% of larvae (Five biological replicates of each genotype with n=30 larvae each). Data presented as a scatter plot with the lines representing the mean and standard deviation. (D-I) Larval intestine from *w^1118^*controls and *D2hgdh* mutants with melanotic masses were dissected and stained with DAPI. (E,F) These lesions are most prevalent in the mutant hindgut and (H,I) are associated with regions that lack of DAPI stained nuclei. (H,I) Red arrows point to melanotic masses in confocal images. The scale bar representing 200 μM in (D) applies to (E,F). The scale bar representing 200 μM in (G) applies to (H,I). All the experiments were repeated three times.

In addition to the melanotic mass phenotype, *D2hgdh^5-5^*and *D2hghd^12-6^* mutant females exhibit reduced fecundity, and embryos laid by mutants displayed reduced viability (Figure 3A,B). However, neither *D2hgdh* mutation affected larval or pupal viability (Figure 3C,D). These phenotypes differ from those reported in previous studies using *UAS-Idh-R195H*, where expression using *Tubulin-Gal4* induced an embryonic lethality, and expression driven by *Ubiquitin-Gal4* and *Actin5C-Gal4* resulted in pupal lethality (REITMAN *et al*. 2015). Thus, developmental phenotypes displayed *D2hgdh* mutants are milder than those induced by *UAS-Idh-R195H* overexpression.

**Figure 3.**
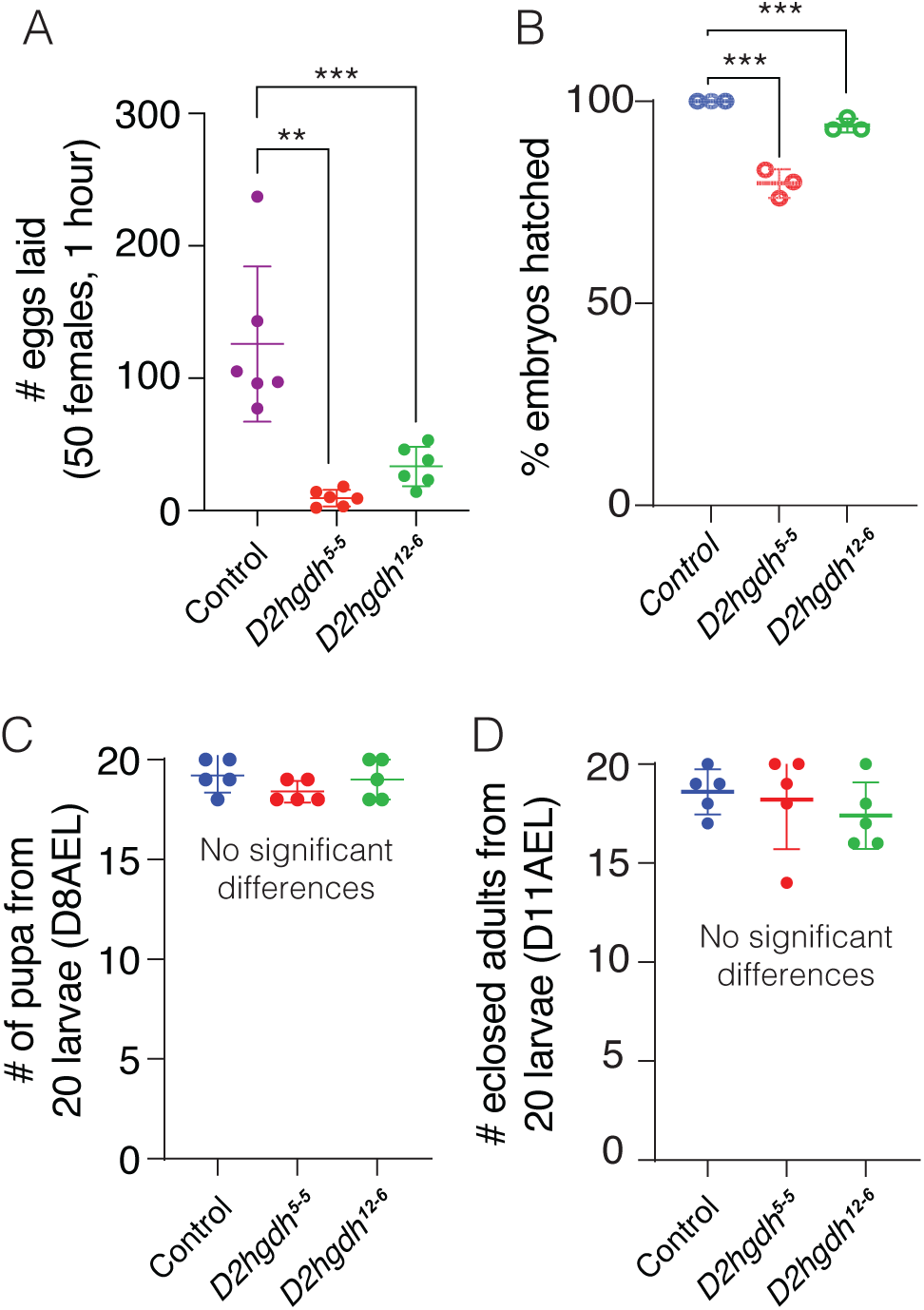
*D2hgdh* mutants exhibit several developmental phenotypes. When compared with the *w^1118^*control strain, both *D2hgdh^12-6^*and *D2hgdh^5-5^* mutants exhibit significant decreases in (A) female fecundity and (B) embryonic viability (assessed at 30 hrs after egg-laying). In contrast, both (C) pupation rate at 8 days after egg-laying (D8AEL) and (D) eclosion rate were similar across all three strains. Data are presented in scatter plots with mean ± standard deviation. *P*-values calculated using a Wilcoxon/Kruskal-Wallis tests followed by a Dunn’s test to compare mutant sample with control. \**P*<0.05. \*\**P*<0.01. \*\*\**P*<0.001. Note that all *P-*values compare the indicated mutant samples with the control. For developmental analysis, n=30 eggs and n=20 larvae were used. All the experiments were repeated three times.

### Metabolomics analysis of *D2hgdh* mutants

To better understand how loss of D2hgdh activity alters *Drosophila* metabolism and physiology, we used a metabolomics approach to compare adult male *D2hgdh^5-5^* and *D2hghd^12-6^* mutants with *w^1118^*controls. Principle component analysis of the resulting datasets revealed that the metabolomic profiles of both mutant strains significantly differ from the control strain but not from each other, demonstrating that loss of D2hgdh activity appreciably alters the metabolome of adult male flies (Figure 4A). Moreover, both mutant strains exhibit similar changes in metabolite abundance (Figure 4B), suggesting that the observed metabolic phenotypes are not the result of genetic background effects.

**Figure 4.**
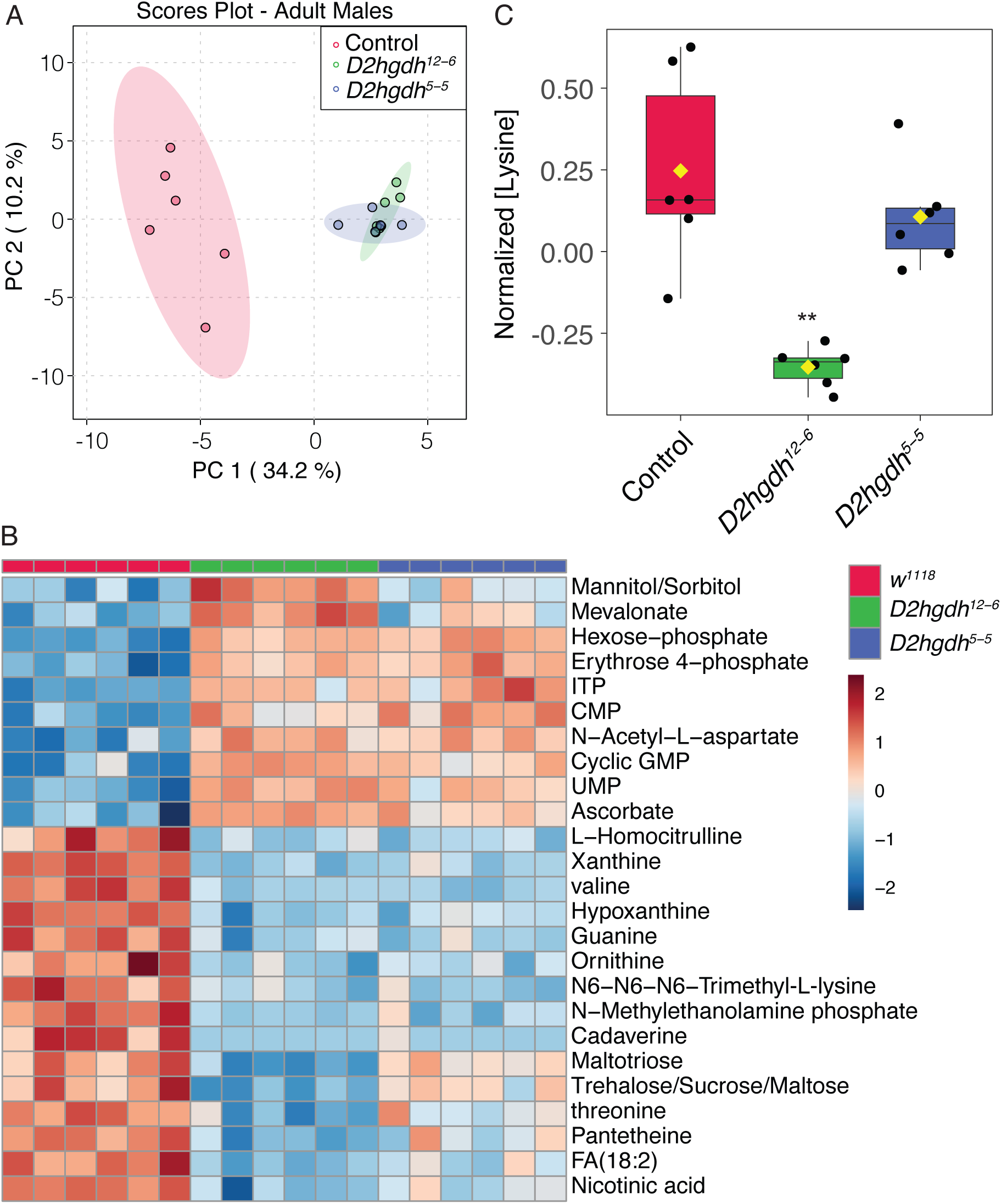
Targeted metabolomic analysis of adult *D2hgdh* mutant males. (A) Adult *D2hgdh* mutant males and *w^1118^*control males were analyzed using a targeted metabolomic approach. (A) Analysis of the metabolomics data using principal component (PC) analysis. Note that the mutant data group together and are entirely separated from the controls. (B) A heatmap illustrating the top 25 most significantly altered compounds in *D2hgdh* mutants as compared with controls. (C) The relative abundance of lysine in adult *D2hgdh* mutant males and a *w^1118^*control strain are presented using a box and whiskers plot. Note that lysine levels are either downregulated or unchanged in mutant strains as compared with the control. Figure panels were generated using Metaboanalyst 5.0, as described in the methods. *P*-values calculated using an ANOVA test followed by a Fisher’s test to compare mutant samples with control. \*\**P*<0.01. A total of six biological replicates were used for each genotype. Each replicate had 20 adults.

KEGG enrichment analysis of both datasets highlighted changes in several metabolic processes, including the pentose phosphate pathway, glutathione, nucleotide, and amino acid metabolism (Figure S1, S2). Consistent with these analyses, both mutant strains exhibit elevated levels of erythrose-4-phosphate, an intermediate in the pentose phosphate pathway, as well as the nucleotides ITP, CTP, UMP, and guanine (Figure 4B). Moreover, we observed decreased levels of the purine degradation products xanthine and hypoxanthine, perhaps suggesting that D-2HG influences purine metabolism. We also observed a highly significant decrease in cadaverine – a breakdown product of lysine commonly found in bacteria. Similarly, N6-N6-N6-trimethyl-L-lysine levels were also significantly lower in mutant males when compared with adults.

Finally, two metabolites, 2HG and lysine, failed to show the expected changes in *D2hgdh* mutants. First, 2HG levels were not significantly altered in our analysis (Table S1). We would note, however, that the semi-targeted analysis used for this study does not differentiate between D-2HG and L-2HG, and thus the observed 2HG values represents the relative abundance of both molecules. As noted above, *D2hgdh* mutant male flies tend to harbor normal levels of L-2HG (See Figure 1), which would partially mask any increase in D-2HG. Second, we also failed to observe an increase in lysine levels, which differs from a recent metabolomic studies of *C. elegans*, where *dhgd-1* mutants displayed increased lysine accumulation (PONOMAROVA *et al*. 2023). In fact, we observed the opposite phenomenon, as lysine was slightly downregulated in one mutant strain as compared with controls (Figure 4C). This result might indicate that *Drosophila* D2hgdh activity influences lysine metabolism in manner that is distinct from that of the *C. elegans* D2hgdh homolog.

## DISCUSSION

Here we establish a new genetic model for studying the role of D2hgdh in animal metabolism, physiology, and development. Our studies demonstrate that D2hgdh is essential for regulating D-2HG levels in the fly, as loss of this enzyme in larvae, adult males, and adult females results in a significant increase in D-2HG levels when compared with a control strain. Moreover, we find that *D2hgdh* mutants display reduced fecundity and embryonic lethality. Finally, metabolomics analysis of these mutants reveals significant changes in the abundance of carbohydrates, nucleotides, and lysine catabolic products. Overall, our studies demonstrate that these *D2hgdh* novel mutations produce a subset of phenotypes consistent with those observed in human D-2-HG aciduria patients and are appropriate for studying D-2HG within the context of human disease models.

While previous studies in *Drosophila* have explored the effects of ectopic D-2HG production using *UAS-Idh-R195H* transgenes (REITMAN *et al*. 2015), the *D2hgdh* loss-of-function mutations described herein provide an opportunity to examine why neomorphic *Idh* mutations and D2hgdh deficiency lead to divergent disease phenotypes despite both inducing aberrant D-2HG accumulation. Interestingly, we find that our *Drosophila D2hgdh* mutants display some phenotypes in common with flies expressing the *UAS-Idh-R195H* transgene. In this regard, both the occasional presence of melanotic masses within *D2hgdh* mutant larvae and the metabolomics profile of adult *D2hgdh* mutant males are notable. The presence of melanotic masses within *D2hgdh* larvae raises several interesting questions about the nature of this phenotype and similarities with human patients. As mentioned above, the human inborn error of metabolism is associated with hypermelanization of the abdominal region (PRESTON *et al*. 2019). While the cause of this phenomenon remains unknown, our observations in the *D2hgdh* mutant, combined with a previous study describing melanotic masses in flies expressing the *UAS-Idh-R195H* transgene in hemocytes, suggest that a conserved metabolic mechanism links elevated D-2HG metabolism with this poorly understood phenomenon. One interesting, albeit speculative, possibility is that the hypermelanization phenotype displayed by *D2hgdh* mutants is that it results from altered immune cell activity – a hypothesis supported by previous studies. First, *hml-Gal4* driven expression of a *UAS-Idh-R195H* transgene in *Drosophila* hemocytes resulted in higher numbers of these cells, demonstrating that D-2HG promotes immune cell development (REITMAN *et al*. 2015). Secondly, excess D-2HG accumulation alters CD8 T lymphocyte activity and *D2hgdh* knockdown decreased the anti-tumor activity of these cells (YANG *et al*. 2022). Future studies should determine if *D2hgdh* mutant larvae display changes in immune cell numbers and/or an altered immune response.

Finally, the metabolomics profile of *D2hgdh* mutants display several unexpected changes, with two trends being of particular interest. First, we observed significant changes in nucleotide metabolism – several nucleotides were significantly upregulated in both mutant background while xanthine and hypoxanthine, products of purine catabolism, are decreased. We are uncertain as to the significance of these findings, however, the results hint at a role for D-2HG in regulating nucleic acid metabolism. The second important observation is that lysine does not accumulate in *D2hgdh* mutants. We expected to observe elevated lysine because (i) the enzyme that degrades lysine is inhibited by D-2HG (CHOWDHURY *et al*. 2011), (ii) *C. elegans dhgd-1* mutants accumulate lysine (PONOMAROVA *et al*. 2023), and (iii) *L2hgdh* mutants exhibit elevated lysine levels (MAHMOUDZADEH *et al*. 2024). Considering that lysine is either unchanged or decreased in *Drosophila* mutants, this result suggests that the relationship between D-2HG and lysine metabolism is more complex than previously anticipated. We would also note lysine is unchanged in D2HG aciduria patients (GIBSON *et al*. 1993; KRANENDIJK *et al*. 2012), thus highlighting a need to further examine this metabolic interaction.

## ACKNOWLEDGEMENTS

We thank the Bloomington *Drosophila* Stock Center (NIH P40OD018537) for providing fly stocks, the *Drosophila* Genomics Resource Center (NIH 2P40OD010949) for genomic reagents, and Flybase (NIH 5U41HG000739). GC-MS was conducted using instruments housed in the Indiana University Mass Spectrometry Facility. A.J.F. was supported by the Indiana University Cox Scholars Program. SS was supported by the John R. and Wendy L. Kindig Fellowship, Briggs Fellowship, Dona Graam Fellowship through Indiana University, and an American Heart Association Predoctoral Fellowship (25PRE1372770). J.M.T. is supported by the National Institute of General Medical Sciences of the National Institutes of Health under a R35 Maximizing Investigators’ Research Award (MIRA; R35GM119557).

## DATA AVAILABILITY

The *w^1118^ D2hgdh^5-5^* and *w^1118^ D2hghd^12-6^* strains have been deposited in the Bloomington *Drosophila* Stock Center and are also available upon request. Metabolomics data is present within Table S1.

## SUPPLEMENTAL FIGURES

**Figure S1. Enrichment analysis of *D2hgdh^5-5^*mutants in comparison to controls.** MetaboAnalyst was used to perform KEGG pathway enrichment analysis of metabolites that were significantly altered in *w^1118^ D2hgdh^5-5^* mutants males as compared with *w^1118^*controls.

**Figure S2. Enrichment analysis of *D2hgdh^12-6^*mutants in comparison to controls.** MetaboAnalyst was used to perform KEGG pathway enrichment analysis of metabolites that were significantly altered in *w^1118^ D2hgdh^12-6^* mutants males as compared with *w^1118^* controls.

## SUPPLEMENTAL TABLE

**Table S1.** Metabolomic analysis of the *D2hgdh* mutants compared with *w^1118^* controls. All samples contained 20 adult males. Data are normalized to sample mass and an internal d4-succininc acid standard.

## Notes

### Competing Interest Statement

The authors have declared no competing interest.

### Summary of Updates

Introduction and Discussion expanded to better compare results in Drosophila with other organisms. Added enrichment analysis of metabolomic data - added two supplemental figures.

