## Supplementary figures and images for "New alleles of *D-2-hydroxyglutarate dehydrogenase* enable studies of oncometabolite function in *Drosophila melanogaster*"

### Figure S1

Control vs *D2hgdh*$^{5\text{-}5}$

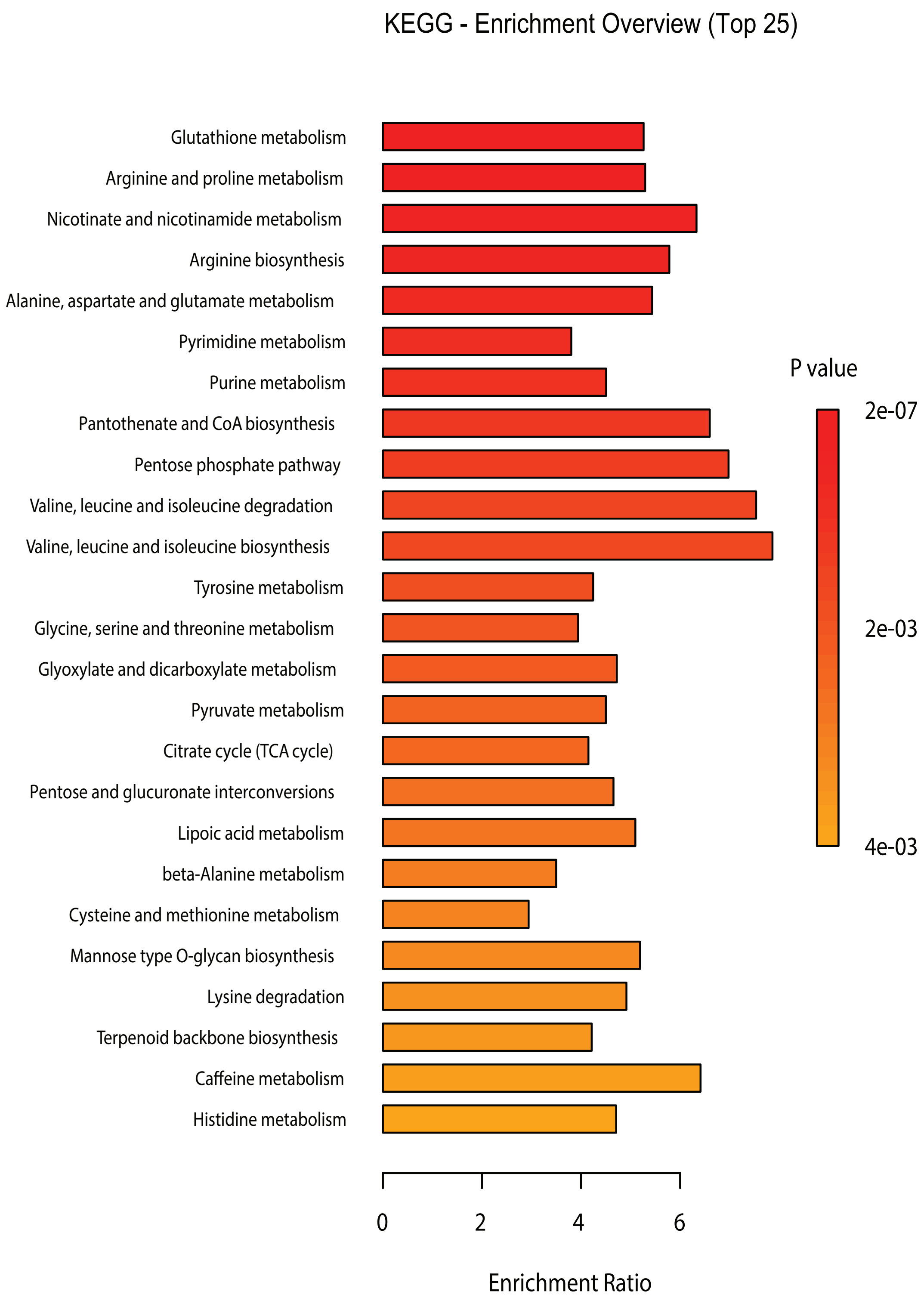


**Figure S2**

### Figure S2

Control vs *D2hgdh*[12-6]

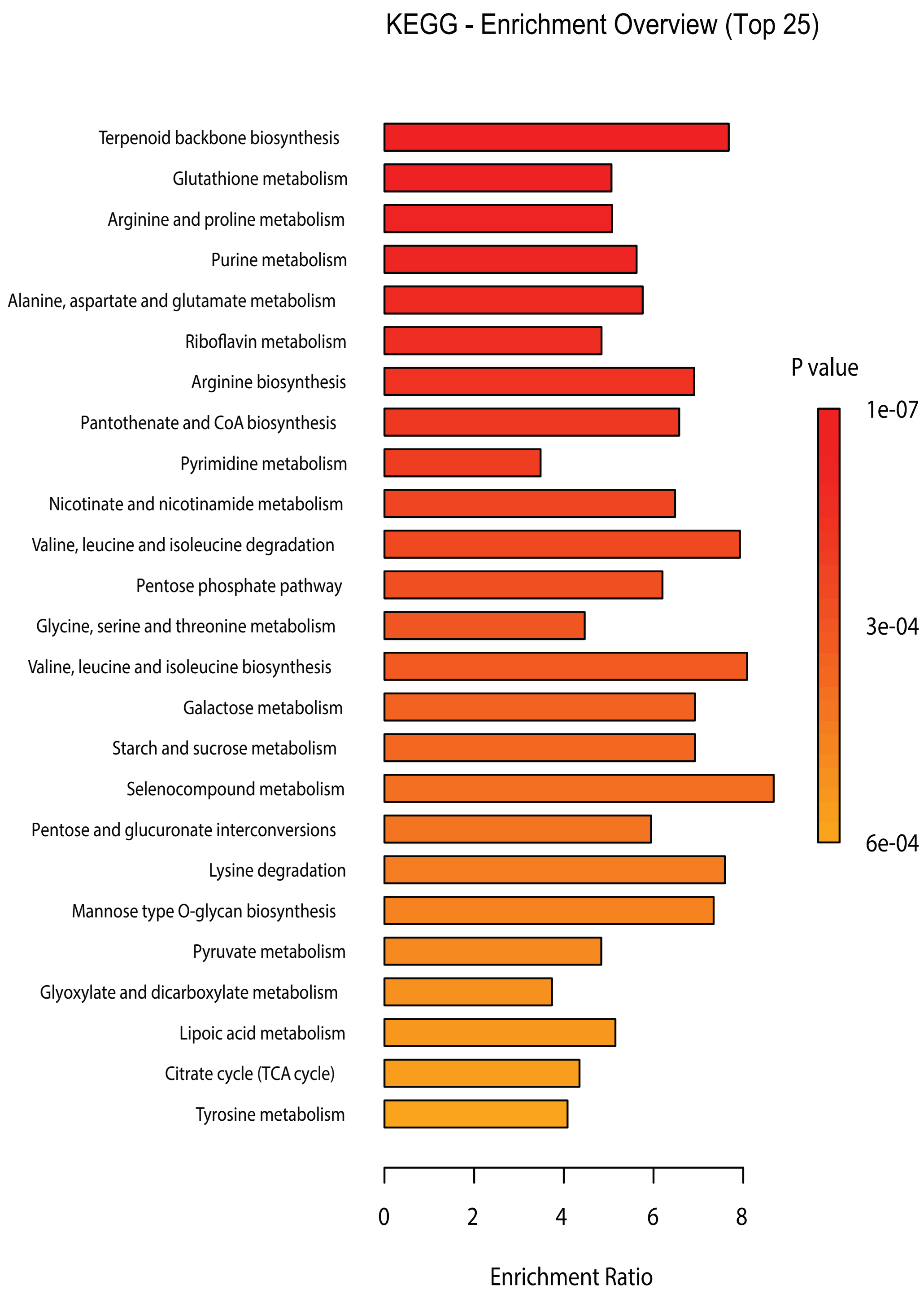


**Figure S3**
